# Evaluation of PicoGreen variants for use in microscopy and flow cytometry

**DOI:** 10.1101/584037

**Authors:** Muznah Khatoo, Jongtae Yang, Kyle R. Gee, Stephan Gasser

## Abstract

PicoGreen is a fluorescent probe that binds dsDNA and forms a highly luminescent complex when compared to the free dye in solution. This unique probe is widely used in DNA quantitation assays but has limited application in flow cytometry and microscopy. Here we have investigated various PicoGreen variants for the ability to stain low amounts of cytosolic DNA present in many tumor cells. Analysis of stained cells by flow cytometry and fluorescent microscopy showed that certain variants improved the ability to stain low levels of cytosolic DNA when compared to the commercially available PicoGreen molecule.

## Introduction

The integrity of DNA is constantly challenged by endogenous and environmental genotoxic factors [1–3]. Cells have evolved elaborate mechanisms to defend the integrity of DNA. Detection of damaged DNA activates complex DNA repair systems and activates innate immune pathways that alert the immune system to the presence of potentially damaged or infected cells [4,5]. In addition, recent studies found that genomic or mitochondrial DNA can accumulate in the cytosol of cells [5]. Similar to damaged DNA, mislocalized DNA triggers innate immune responses including the production of type-I interferons and other inflammatory cytokines [6]. The detection of DNA in the cytosol requires sensitive tools that can be used in conjunction with mitochondrial, nuclear and membrane markers in microscopy and flow cytometry.

Fluorescent probes that specifically bind nucleic acids are useful tools to study DNA metabolism [7]. PicoGreen (PG) is a highly sensitive fluorescent dye developed in the 1990s that detects double-stranded (ds) DNA [8,9]. PG shows significant selectivity for dsDNA and DNA:RNA hybrids over single-stranded (ss) DNA and RNA. Binding to linear dsDNAs results in slightly higher signals when compared to supercoiled plasmids. PG is able to strongly bind to highly polymeric DNA and short DNA duplexes <20 bp. Binding of dsDNA by PG is not affected by the presence of salts, proteins, poly(ethylene glycol), urea, chloroform, ethanol, and agarose, while some ionic detergents and heparin interfere with PG binding. PG has an excitation peak at 480 nm and an emission peak at 520 nm. Upon binding DNA, PG fluorescence increases >1000-fold while unbound PG has virtually no fluorescence. PG is very stable to photo-bleaching, which allows longer exposure times.

## Material and Methods

### Cell lines

Cervical adenocarcinoma HeLa cells were purchased from ATCC. The TRAMP-C2 prostate cancer cell line was kindly provided by Dr. D. H. Raulet (University of California, Berkeley, USA). TRAMP-C2 cells were derived from the transgenic adenocarcinoma mouse prostate (TRAMP) model which specifically expresses SV40 large T-antigen in prostate epithelial cells [10]. TRAMP-C2 and HeLa cells were cultured in Dulbecco’s Modified Eagle Medium (DMEM) medium (Invitrogen, ThermoFisher) supplemented with 10% fetal calf serum (FCS; Hyclone, GE Healthcare), 50 μM 2-mercaptoethanol, 200 μM asparagine, 2 mM glutamine (Sigma, Merck) and 1% penicillin-streptomycin (Pen-Strep; Invitrogen, ThermoFisher). All cells were treated with Plasmocin (Invivogen) to exclude potential mycoplasma contaminations.

### Reagents

PicoGreen, OliGreen, SlowFade Gold Antifade, ProLong Gold Antifade, Image-iT FX Signal Enhancer, lambda DNA, ssDNA (M13 primer ssDNA-M13 primer 5’-TGT-AAA-ACG-ACG-GCC-AGT-3’, paraformaldehyde (PFA) were purchased from ThermoFisher. PicoGreen variants were synthesized in house at ThermoFisher. Lipofectamine 3000 was used for transfection of dsDNA and ssDNA according to the manufacturer’s protocol. 10 ug of lamda DNA (NEB) or ssDNA M13 primer was labelled using the Ulysis Alexa Fluor 647 nucleic acid labeling kit (Thermo Fisher) according to the manufacturer’s instructions.

### Microscopy

Cells were stained with 3 μl/ml of PicoGreen variants (200 μM) or OliGreen for 90 min at 37°C. For some experiments, cells were fixed in 4% PFA for 10 min at room temperature prior to PicoGreen staining. Stainings were analyzed using an Olympus FV1000 confocal scanning microscope equipped with a 100 x oil immersion objective and an ApoTome optical sectioning device (Zeiss). Pictures were analyzed using Photoshop CS5 (Adobe, USA), Metamorph NX (Molecular Devices) or Imaris X64 (Version 7.6.4, Bitplane). Imaris software was used to calculate the Pearson’s correlation coefficient.

### Flow Cytometry

Cells were stained as described above with PicoGreen or OliGreen. Stained cells were analyzed by multicolor flow cytometry using FACSCalibur (BD Biosciences, USA) or LSRFortessa (BD Biosciences) and FlowJo 8.8.6 software (Treestar, USA).

### Statistical Analyses

Statistical analyses were performed using student’s t-test. Error bars denote ±SD. \**p*<0.05; \*\**p*<0.01; \*\*\**p*<0.001.

## Results

### Enhanced stainings of DNA by PicoGreen variants in confocal microscopy analysis

We have recently shown the presence of cytosolic DNA in many human cancers and cancer cell lines including TRAMP-C2 and HeLa cells [11–13]. To evaluate the ability of asymmetrical cyanine dyes to label cytosolic DNA, we stained the murine prostate cell line TRAMP-C2 with 65 different PicoGreen variants, all of which share the same core structure (Fig 1). Analysis of live cells staining of TRAMP-C2 cells identified a number of PicoGreen variants that stained cytosolic DNA with higher intensity than the commercially available PicoGreen. The brighter cytosolic DNA signals by PicoGreen variants #2, #34, #41 and #48 correlated with enhanced nuclear DNA staining suggesting that these variants result in stronger fluorescent signals upon binding of dsDNA (Fig 1).

**Fig 1.**
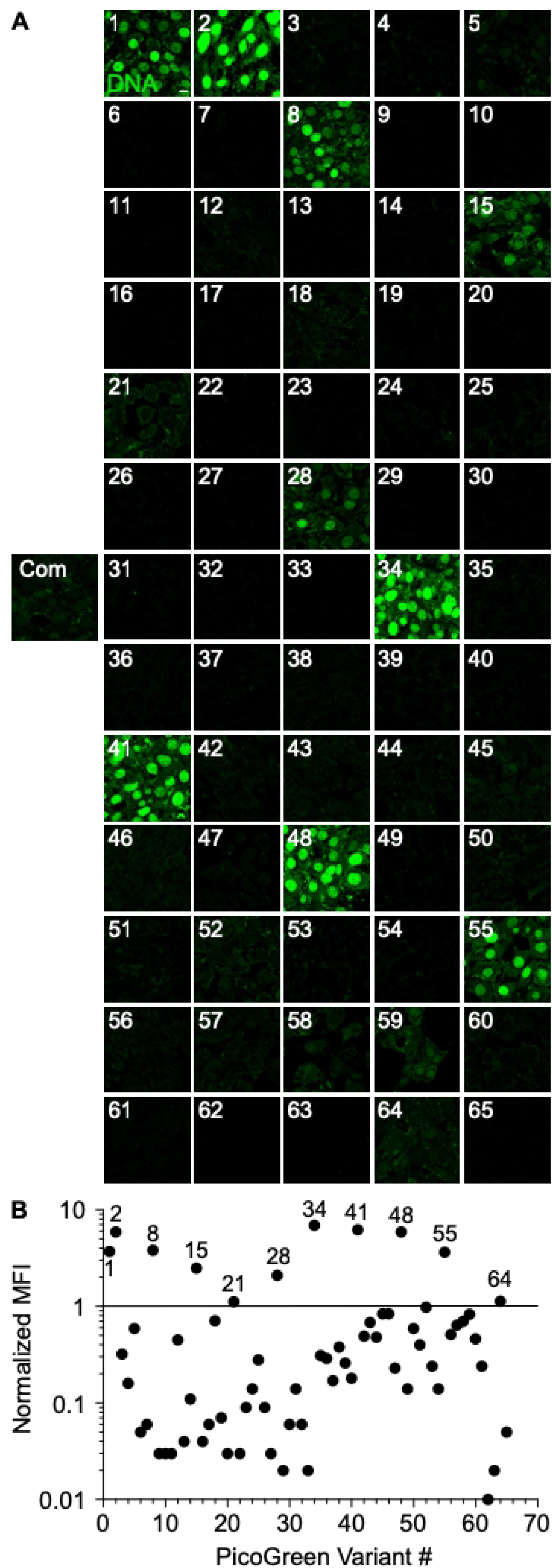
Confocal analysis of TRAMP-C2 cell staining with PicoGreen variants. (A) Confocal images of 4% paraformaldehyde-fixed TRAMP-C2 cells stained with 3 μl/ml of commercially available PicoGreen variants (Green) or 65 different PicoGreen variants (green) for 90 min. (B) Quantification of PicoGreen mean fluorescence intensity (MFI) normalized to commercially available PicoGreen. Scale bar denotes 10 μm.

### Enhanced stainings of DNA by PicoGreen variants in flow cytometric analysis

Next, we tested the ability of PicoGreen variants to stain dsDNA present in TRAMP-C2 cells by flow cytometry. Similar to our findings using microscopy to analyze the cytoplasmic PicoGreen signals, staining of dsDNA with the PicoGreen variants #34, #41 and #48 resulted in the highest mean fluorescence (MFI) when compared to the commercially available PicoGreen (Fig 2). Interestingly, the MFI of PicoGreen variant #2, which was among the best variants for staining of dsDNA in microscopy, was lower than the commercially available PicoGreen dye suggesting that PicoGreen variant #2 is more sensitive to photobleaching by the more powerful 488 nm laser used in flow cytometry.

**Fig 2.**
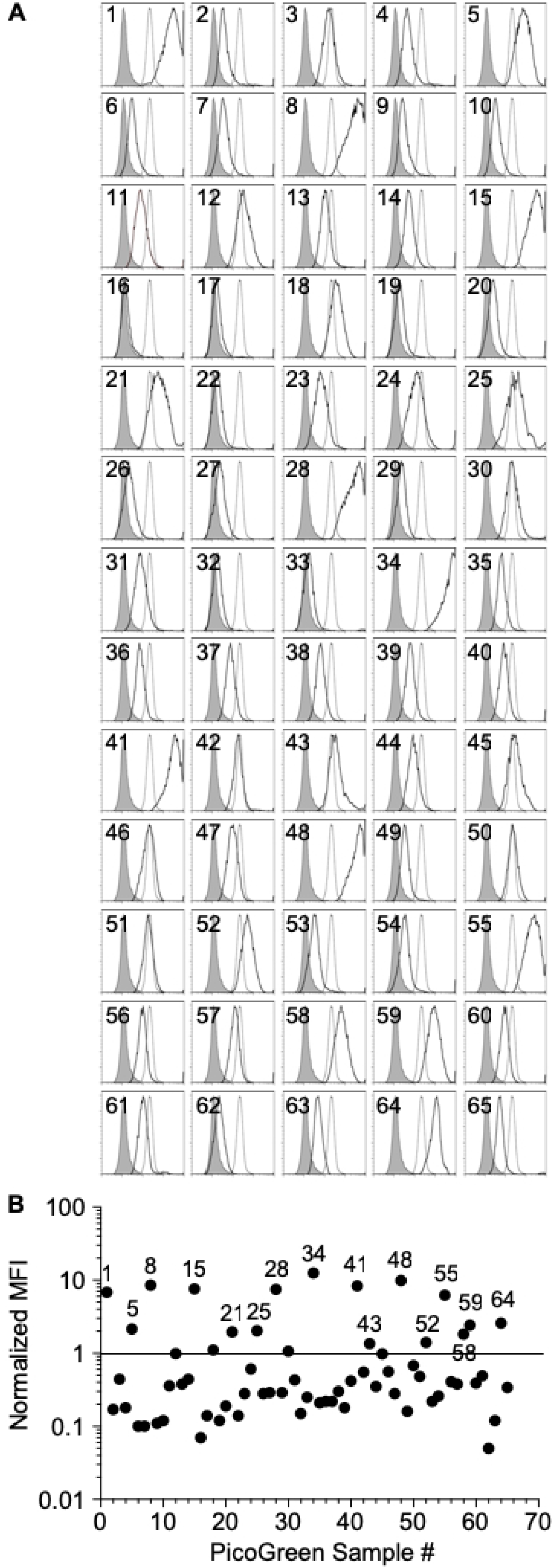
Flow cytometric analysis of TRAMP-C2 cell staining with PicoGreen variants. (A) Flow cytometric analysis of 4% paraformaldehyde-fixed TRAMP-C2 cells stained with 3 μl/ml of commercially available PicoGreen variants (Green) or 65 different PicoGreen variants (green) for 90 min. (B) Quantification of PicoGreen mean fluorescence intensity (MFI) normalized to commercially available PicoGreen.

### Fixation condition impact staining of DNA by PicoGreen variants

In microscopy, fixation is often a prerequisite to further processing of clinical samples. For that reason, we tested the ability of PicoGreen variants to stain dsDNA in paraffin- and paraformaldehyde (PFA)-fixed TRAMP-C2 cells. The PicoGreen variants with the highest normalized MFI (Fig 1) for staining of dsDNA in live cells were chosen. Paraffin-fixed TRAMP-C2 cells were smaller in size and no clear cytosolic dsDNA signal could be detected for any of the PicoGreen variants (Fig 3). In contrast, fixation of cells with 4% PFA slightly enhanced the staining intensity of PicoGreen and also increased the resolution of the cytosolic DNA staining. However, PFA fixation abrogated the ability of the PicoGreen variant #55 to stain cytosolic DNA (Figs 1 and 3). In summary, our data suggest that PFA fixation and the use of certain PicoGreen variants can drastically increase dsDNA staining signals and resolution.

**Fig 3.**
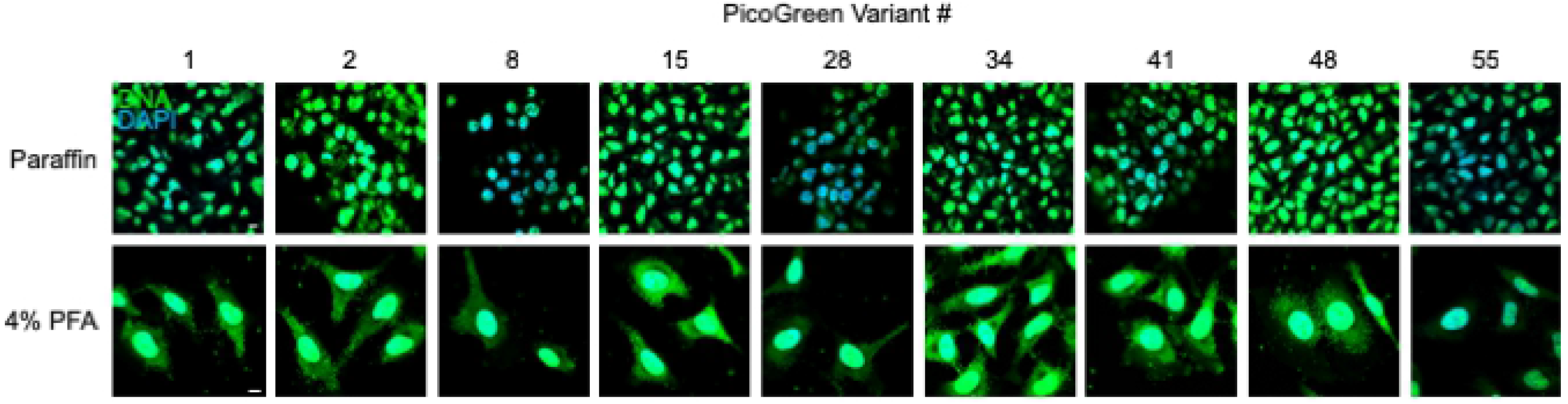
Confocal analysis of paraffin-fixed TRAMP-C2 cells stained with PicoGreen variants. Confocal images of paraffin-fixed (upper row) or 4% paraformaldehyde-fixed (lower row) TRAMP-C2 cells stained with 3 μl/ml of commercially available PicoGreen variants (Green) or PicoGreen variants #1, #2, #8, #15, #28, #34, #41, #48 or # 55 (green) for 90 min in the presence of DAPI (blue) to reveal nuclear DNA. Scale bar denotes 10 μm.

### Anti-fade reagents fail to enhance the staining of PicoGreen variants

Anti-fade reagents were shown to suppress photobleaching and cause little or no quenching of the initial fluorescent signal [14,15]. Slowfade and Slowfade Gold are glycerol-based mountants of several commonly used fluorophores, but both reagents failed to enhance PicoGreen staining of cytosolic DNA (Fig 4). Slowfade Gold quenched the fluorescence of PicoGreen variant #15 and #28. A similar trend was observed for the liquid mountant Prolong Gold. Finally, no increased cytosolic DNA staining was observed by the different PicoGreen variants in presence of Image-IT Fx Signal enhancer, which can block background staining that results from non-specific interactions of a wide variety of fluorescent dyes with cell and tissue constituents. In summary, several anti-fade reagents were not able to enhance the signal to noise ratio of cytosolic DNA staining by different PicoGreen variants.

**Fig 4.**
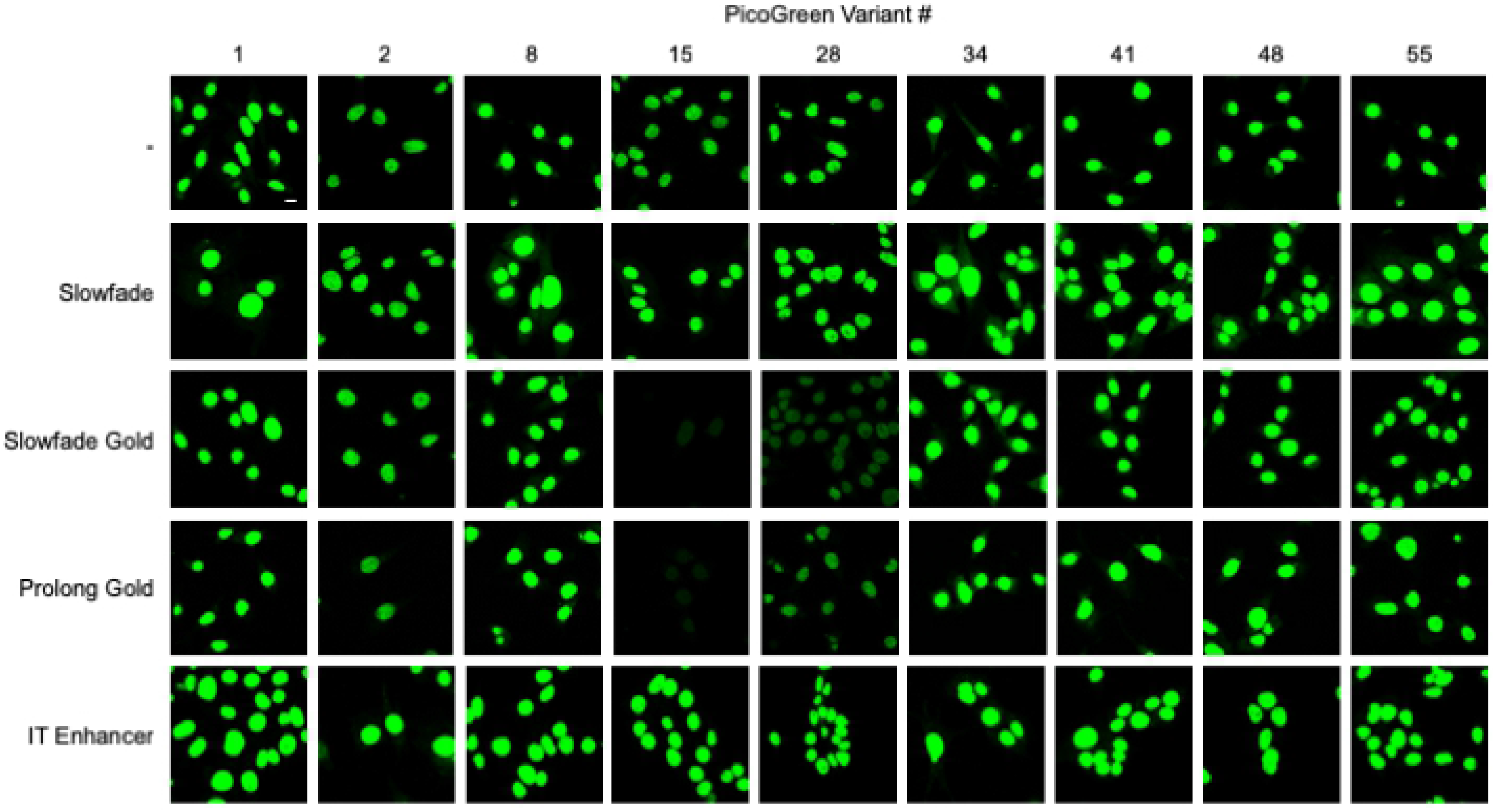
Confocal analysis of the effect of anti-fade agents on TRAMP-C2 cell staining with PicoGreen variants. Confocal images of 4% paraformaldehyde-fixed TRAMP-C2 cells stained with 3 μl/ml of commercially available PicoGreen variants or PicoGreen variants #1, #2, #8, #15, #28, #34, #41, #48 or # 55 (green) for 90 min in the presence of SlowFade, SlowFade Gold, ProLong Gold antifade mountant or IT enhancer. Scale bar denotes 10 μm.

### PicoGreen variants specifically recognize dsDNA

To analyze the specificity of the different PicoGreen variants towards different nucleic acid types, we measured the fluorescence enhancement *in vitro* by spectroscopy when the different dyes bound to dsDNA, ssDNA, ssRNA or dsRNA (Fig 5). All variants showed specificity towards dsDNA when compared to ssDNA, ssRNA and dsRNA. Some dyes such as PicoGreen variant #34, #41, #48 and #55 also weakly bound to dsRNA and to a lesser degree ssDNA. In summary, the PicoGreen variants #34, #41 and #48 showed the greatest fluorescence enhancement when bound to 500 ng/ml dsDNA, which was more than 2-fold higher than the commercially available PicoGreen (Fig 5A). These results are in agreement with our earlier observation that the PicoGreen variants #34, #41 and #48 resulted in the highest fluorescence when analyzed by microscopy (Fig 1) or flow cytometry (Fig 2). The differences in fluorescence enhancement were less pronounced when smaller amounts of dsDNA were measured (Figs 5B and 5C), but most PicoGreen variants still outperformed the commercially available PicoGreen.

**Fig 5.**
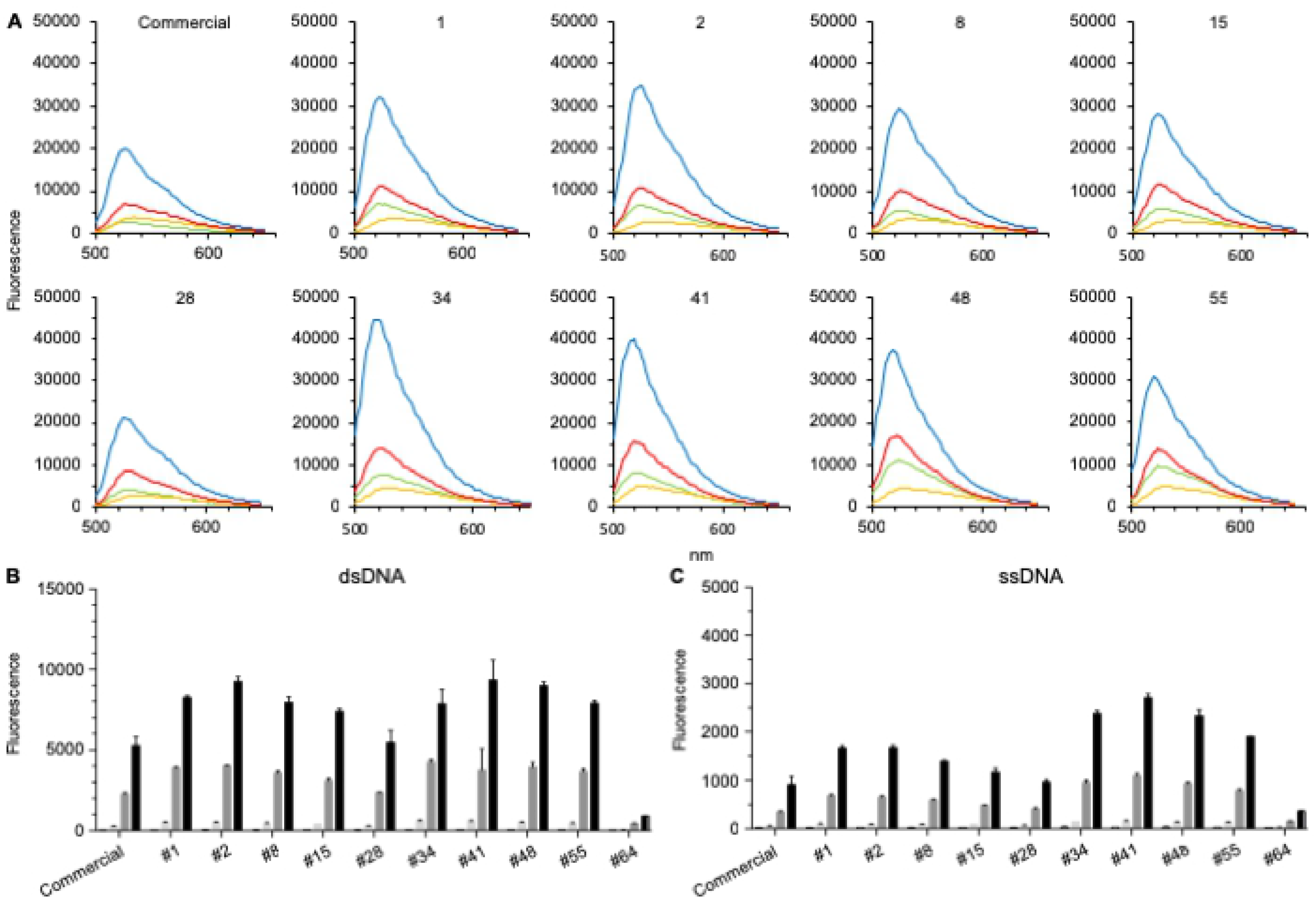
*In vitro* dsDNA specificity of PicoGreen variants. (A) 3 μl/ml of commercial PicoGreen or PicoGreen variants were added to 500 ng/mL dsDNA (blue), ssDNA (green), RNA (yellow) or microRNA (red). (B-C) 3 μl/ml of commercial PicoGreen or PicoGreen variants were added to 0 ng/ml (white columns), 0.5 ng/ml (light grey columns), 4 ng/ml (dark grey columns) or 10 ng/ml (black columns) dsDNA (B) or ssDNA (C). Samples were read on a Tecan plate reader at 491 nm excitation and 500-650 nm (A) or 520 nm (B, C) emission. The mean and standard deviation of triplicates is shown.

### In vivo specificity of PicoGreen variants

To examine the specificity to the PicoGreen variants towards dsDNA in cells, we stained HeLa cells transfected with Alexa647-labelled dsDNA or ssDNA. The PicoGreen variants #41 and #48 stained cytosolic DNA and a majority of the transfected labelled dsDNA (Fig 6). In contrast, commercially available PicoGreen or the PicoGreen variants failed to stain transfected dsDNA, which correlated with weaker staining of endogenous DNA. The PicoGreen variants #41 and #48 also stained some of the transfected Alexa647-labelled ssDNA consistent with a weak binding to ssDNA in vitro. It is possible that high local concentration of ssDNA and/or formation of dsDNA due to interaction with cytosolic DNA enabled sufficient enhancement of fluorescence by the PicoGreen variants #41 and #48. In summary, PicoGreen variants #41 and #48 show enhanced staining of cytosolic dsDNA when compared to commercially available PicoGreen. These PicoGreen variants might be useful to better understand the nature and behaviour of cytosolic DNA in cancer or infected cells.

**Fig 6.**
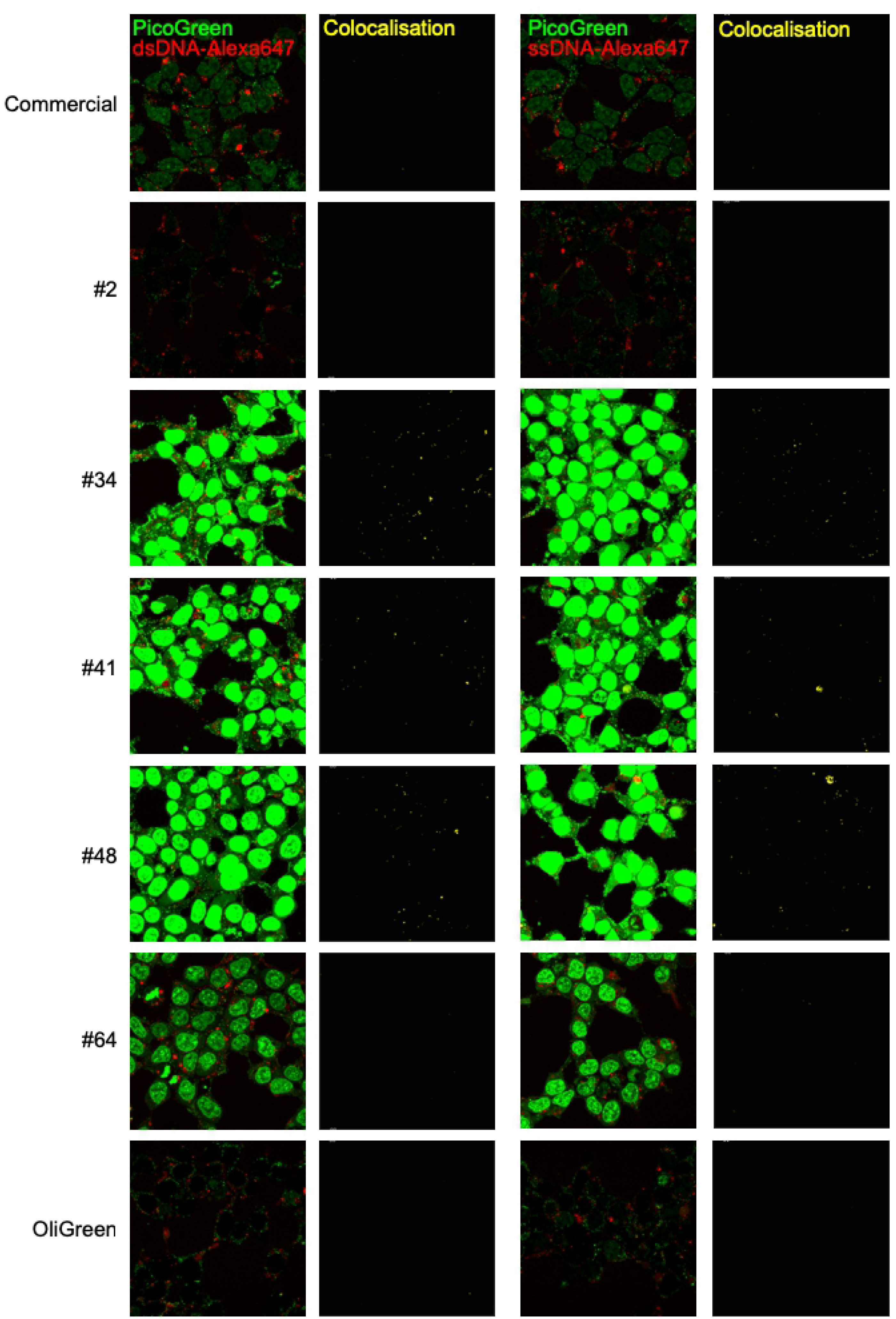
*In vivo* dsDNA specificity of PicoGreen variants. 293T cells were transfected with 1 μg of Alexa647-labelled dsDNA (left columns) or Alexa647-labelled ssDNA. Transfected cells were stained with 3 μl/ml of commercially available PicoGreen variants or PicoGreen variants #1, #2, #8, #15, #28, #34, #41, #48 or # 55 (green) for 90 min. Co-localized regions are shown in yellow in the right panels. Co-localization was calculated using the PCC equation in Imaris for Alexa647^+^ cells including nuclear PicoGreen signals. Scale bar denotes 10 μm.

## Discussion

To visualize DNA present in the cytosol of many tumor cells, we have previously used commercially available PicoGreen [11–13]. The fluorescence enhancement of PicoGreen when bound to dsDNA is very high with negligible background signal from unbound PicoGreen molecules [16]. PicoGreen fluorescence signals remain stable during the image acquisition as it is relatively insensitive to photobleaching. We have successfully used PicoGreen for up to 90 minutes in live-cell imaging [11]. PicoGreen stainings of tumor cells suggested a granular distribution of cytosolic DNA in tumor cells, while stainings using dsDNA-specific antibodies indicated a more homogenous distribution of DNA in the cytosol possibly in part due to greater sensitivity of the antibody staining [11]. PicoGreen was initially developed for quantification of DNA by spectrofluorometer [16]. We therefore reevaluated 65 PicoGreen variants of the original variant library for their ability to stain cytosolic DNA by flow cytometry and fluorescence microscopy with greater sensitivity than the commercially available PicoGreen. Screening of 65 PicoGreen variants showed that the variants #2, #34, #41 and #48 correlated with enhanced staining of cytosolic and nuclear DNA. The enhanced cytosolic DNA signals is likely due to the stronger emission upon binding of dsDNA by the variants #2, #34, #41 and #48 when compared to commercially available PicoGreen. Importantly, the PicoGreen variants #2, #34, #41 and #48 maintain specificity towards dsDNA as the low fluorescence upon dsRNA, ssRNA or ssDNA binding was similar to the commercially available PicoGreen.

In agreement with the dsDNA-specific antibody stainings of cytosolic DNA in tumor cells, the brighter PicoGreen variants revealed additional cytosolic DNA in the cytosol not observed with commercially available PicoGreen. Interestingly, the staining of DNA in the cytosol with the brighter PicoGreen variants indicated differential distribution of DNA some of which showed granular distribution, while other cytosolic DNA appeared to be more homogenously distributed in the cytosol of tumor cells. It is likely that the granular staining of DNA is reflecting the aggregation of dsDNA in the cytosol as a similar granular accumulation of dsDNA was observed upon staining with dsDNA-specific antibodies ([11] and data not shown). We have previously shown that granular DNA staining in the cytoplasm does not overlap with dsDNA present in mitochondria, but it possible that mitochondrial DNA contributes to the DNA staining outside the granules [11]. Furthermore, it is possible that other nucleotide species such as RNA:DNA hybrids [11–13] and long dsRNAs [17] are stained by the PicoGreen variants.

The increased fluorescence emission of dsDNA-bound PicoGreen depends on the structure and binding mode of PicoGreen [9,18]. PicoGreen has a 4-[[2,3-dihydro-3-methyl-(benzo-1,3-thiazol-2-yl)-methylidene]-quinolini-um]+ core structure. In addition, it contains a phenyl group at position one of the quinolinium ring and a N-bis-(3-dimethylaminopropyl)-amino residue at position two of the quinolinium ring. PicoGreen carries three positive charges, which are likely to contribute to its high binding affinity for dsDNA. We observed that electron donating substituents on the delocalized benzothiazolyl-quinolinium system common to PicoGreen variants correlate with brighter signals from cellular dsDNA. Electron-donating groups were previously found to form more stable ligand-DNA complexes [19]. In addition, the electron donating substituents can also induce an increase in the molar absorption coefficient and a shift in the fluorescence spectra, which was not observed for the different PicoGreen variants analyzed in this study (Fig 5) [20].

Photobleaching is an irreversible photochemical process that stops the emission of photons by fluorophores [21]. The bleaching process is likely to involve reaction of excited-dye molecules with reactive oxygen species (ROS) that are produced when molecular oxygen interacts with different electronic states of the dye [22]. Most antifade reagents are reactive oxygen species scavengers. Glycerol-based Prolong was reported to have among the highest anti-fading properties for many applications while good antifading properties have also been reported for the other reagents used in this study [23,24]. Surprisingly, none of the anti-fading reagents including Prolong improved the staining of cytosolic DNA in tumor cells. In fact, Prolong and Slowfade Gold decreased the staining intensity of some PicoGreen variants. Consistent with this finding, it was previously found that antifade reagents can sometimes negatively affect fluorophores by quenching the fluorescence of the dye [14].

In summary, we show in this study that the addition of electron donating substituents to the commercially PicoGreen enhances the staining of DNA in confocal microscope and flow cytometry. The high sensitivity and specificity of PicoGreen for dsDNA in addition to the ease of use should enable wide spread use of PicoGreen variants in many applications.

## Author contributions

M.K. performed the experiments and analysed the data. J.Y. performed and analysed the data in figure 5. K.R.G. and S.G. analysed all data, designed experimental strategies and wrote the paper.

## Conflict of interest

The authors declare no conflict of financial interests.

## Acknowledgments

We would like to thank Ms S.Y. Lee for her help in imaging cells. This work was supported by the ThermoFisher Discovery Innovation Grant RL2015-137-(LTC-NUS).

## References

1. Lindahl T. Instability and decay of the primary structure of DNA. Nature. 1993;362: 709–715. doi:10.1038/362709a0

2. Kunkel TA. DNA replication fidelity. J Biol Chem. 2004;279: 16895–16898. doi:10.1074/jbc.R400006200

3. Nyberg KA, Michelson RJ, Putnam CW, Weinert TA. Toward Maintaining the Genome: DNA Damage and Replication Checkpoints. Annu Rev Genet. 2002;36: 617–656. doi:10.1146/annurev.genet.36.060402.113540

4. Goldstein M, Kastan MB. The DNA damage response: implications for tumor responses to radiation and chemotherapy. Annu Rev Med. 2015;66: 129–143. doi:10.1146/annurev-med-081313-121208

5. Gasser S, Zhang WYL, Tan NYJ, Tripathi S, Suter MA, Chew ZH, et al. Sensing of dangerous DNA. Mechanisms of Ageing and Development. 2016. doi:10.1016/j.mad.2016.09.001

6. Wu J, Chen ZJ. Innate Immune Sensing and Signaling of Cytosolic Nucleic Acids. Annu Rev Immunol. 2014;32: 461–488. doi:10.1146/annurev-immunol-032713-120156

7. Boutorine AS, Novopashina DS, Krasheninina OA, Nozeret K, Venyaminova AG. Fluorescent probes for nucleic Acid visualization in fixed and live cells. Molecules. 2013;18: 15357–15397. doi:10.3390/molecules181215357

8. Dragan AI, Casas-Finet JR, Bishop ES, Strouse RJ, Schenerman MA, Geddes CD. Characterization of PicoGreen interaction with dsDNA and the origin of its fluorescence enhancement upon binding. Biophys J. 2010;99: 3010–3019. doi:10.1016/j.bpj.2010.09.012

9. Zipper H, Brunner H, Bernhagen J, Vitzthum F. Investigations on DNA intercalation and surface binding by SYBR Green I, its structure determination and methodological implications. Nucleic Acids Res. 2004;32: e103–e103. doi:10.1093/nar/gnh101

10. Hurwitz A, Foster B, Allison J, Greenberg N, Kwon E. The TRAMP mouse as a model for prostate cancer. Curr Protoc Immunol. 2001;Chapter 20: Unit 20 5. doi:10.1002/0471142735.im2005s45

11. Shen YJ, Le Bert N, Chitre AA, Koo CX, Nga XH, Ho SSW, et al. Genome-Derived Cytosolic DNA Mediates Type I Interferon-Dependent Rejection of B Cell Lymphoma Cells. Cell Rep. 2015;11: 460–473. doi:10.1016/j.celrep.2015.03.041

12. Lam AR, Le Bert N, Ho SSW, Shen YJ, Tang LFM, Xiong GM, et al. RAE-1 ligands for the NKG2D receptor are regulated by STING-dependent DNA sensor pathways in lymphoma. Cancer Res. 2014. doi:10.1158/0008-5472.CAN-13-1703

13. Ho SSW, Zhang WYL, Tan NYJ, Khatoo M, Suter MA, Tripathi S, et al. The DNA Structure-Specific Endonuclease MUS81 Mediates DNA Sensor STING-Dependent Host Rejection of Prostate Cancer Cells. Immunity. 2016;44: 1177–1189. doi:10.1016/j.immuni.2016.04.010

14. Longin A, Souchier C, Ffrench M, Bryon PA. Comparison of anti-fading agents used in fluorescence microscopy: image analysis and laser confocal microscopy study. J Histochem Cytochem. 2017;41: 1833–1840.

15. Albrecht RM, Oliver JA. Labeling Considerations for Confocal Microscopy. Basic Confocal Microscopy. New York, NY: Springer New York; 2011. pp. 79–114.

16. Singer VL, Jones LJ, Yue ST, Haugland RP. Characterization of PicoGreen reagent and development of a fluorescence-based solution assay for double-stranded DNA quantitation. Anal Biochem. 1997;249: 228–238. doi:10.1006/abio.1997.2177

17. Nguyen TA, Smith BRC, Tate MD, Belz GT, Barrios MH, Elgass KD, et al. SIDT2 Transports Extracellular dsRNA into the Cytoplasm for Innate Immune Recognition. Immunity. 2017;47: 498–509.e6. doi:10.1016/j.immuni.2017.08.007

18. Wiederschain GY. The Molecular Probes handbook. A guide to fluorescent probes and labeling technologies. Biochemistry Moscow. SP MAIK Nauka/Interperiodica; 2011;76: 1276–1276. doi:10.1134/S0006297911110101

19. Jain AK, Gupta SK, Tandon V. Evaluation of electronic effect of phenyl ring substituents on the DNA minor groove binding properties of novel bis and terbenzimidazoles: synthesis and spectroscopic studies of ligand-DNA interaction. Oligonucleotides. 2009;19: 329–340. doi:10.1089/oli.2009.0190

20. Valeur B, Berberan-Santos MN. Molecular fluorescence: principles and applications. 2012.

21. Lichtman JW, Conchello J-A. Fluorescence microscopy. Nat Methods. 2005;2: 910–919. doi:10.1038/nmeth817

22. Stennett EMS, Ciuba MA, Levitus M. Photophysical processes in single molecule organic fluorescent probes. Chem Soc Rev. 2014;43: 1057–1075. doi:10.1039/c3cs60211g

23. Ono M, Murakami T, Kudo A, Isshiki M, Sawada H, Segawa A. Quantitative comparison of anti-fading mounting media for confocal laser scanning microscopy. J Histochem Cytochem. 2001;49: 305–312. doi:10.1177/002215540104900304

24. Longin A, Souchier C, Ffrench M, Bryon PA. Comparison of anti-fading agents used in fluorescence microscopy: image analysis and laser confocal microscopy study. J Histochem Cytochem. 1993;41: 1833–1840.

